# Impact of South African 501.V2 Variant on SARS-CoV-2 Spike Infectivity and Neutralization: A Structure-based Computational Assessment

**DOI:** 10.1101/2021.01.10.426143

**Authors:** Mary Hongying Cheng, James M Krieger, Burak Kaynak, Moshe Arditi, Ivet Bahar

## Abstract

**Motivation:** The SARS-CoV-2 variants emerging from South Africa (501.V2) and the UK (B.1.1.7) necessitate rapid assessment of the effects of the corresponding amino acid substitutions in the spike (S) receptor-binding domain (RBD) of the variants on the interactions with the human ACE2 receptor and monoclonal antibodies (mAbs) reported earlier to neutralize the spike.

**Results:** Molecular modeling and simulations reveal that N501Y, shared by both variants, increases ACE2 binding affinity, and may impact the collective dynamics of the ACE2-RBD complex, occupying a central hinge site that modulates the overall dynamics of the complex. In contrast, the substitutions K417N and E484K in the South African variant 501.V2 would reduce the ACE2-binding affinity by abolishing two interfacial salt bridges that facilitate RBD binding to ACE2, K417(S)-D30(ACE2) and E484 (S)-K31(ACE2). These two mutations may thus be more than compensating the attractive effect induced by N501Y, overall resulting in an ACE2-binding affinity comparable to that of the wildtype RBD. Further analysis of the impact of these mutations on the interactions with mAbs targeting the spike indicate that the substitutions K417N and E484K may also abolish the salt bridges between the spike and selected mAbs, such as REGN10933, BD23, H11_H4, and C105, thus reducing the binding affinity and effectiveness of these mAbs.

**Supplementary information:** Supplementary data are available at *Bioinformatics* online.

## 1 Introduction

The high transmissibility of SARS-CoV-2 and the lack of a robust pre-existing or acquired immunity by the hosts (Petersen, et al., 2020; Tortorici, et al., 2020) have caused COVID-19 cases to surge to more than 83 million over the world, resulting in over 1.8 million deaths by the end of 2020. There is an urgent need to prevent infection through vaccination and/or develop effective therapies either by monoclonal antibody (mAb) treatments or development of antiviral compounds development to prevent and treat COVID-19 and associated multi-system inflammatory syndromes. Two vaccines produced by Pfizer/BioNTech and Moderna have already been authorized for emergency use in the US in December 2020, while the duration of the immune protection, their ability to protect populations across all age groups, as well as the capacity to produce sufficient doses of safe and effective vaccines remain to be determined (Krammer, 2020). Most importantly, even though the mutation rate of SARS-CoV-2 has been slower than that of SARS-CoV of 2003 and other SARS-family coronaviruses, new SARS-CoV-2 variants emerged in recent weeks, which apparently increase transmissibility and induce immune escape (Andreano, et al., 2020; Kemp, et al., 2020; McCarthy, et al., 2020). Open genomic surveillance data sharing and collaborative online platforms allow us a real-time tracking of the emergence and spread of many virus lineages (Hadfield, et al., 2018; Shu and McCauley, 2017). As of December 11^th^, 2020, 246,534 SARS-CoV-2 sequences have been listed in the GIASID database (https:gisaid.org). Of note are the so-called UK B1.1.7 (Kemp, et al., 2020) and Southern African 501.V2 variants (Tegally, et al., 2020) which appear to be highly transmissible and have now been observed in multiple countries.

The SARS-CoV-2 spike (S) glycoprotein plays a major role in infectivity. S is a trimer, each monomer being composed of two subunits, S1 and S2. The receptor binding domain (RBDs; residues 331-524) of the S1 subunit recognizes the human receptor angiotensin-converting enzyme 2 (ACE2); and the S2 trimer enables cell fusion and/or endocytosis following two cleavages by human proteases and ensuing massive conformational change. Not surprisingly, the majority of COVID-19 mAb therapies under investigation are designed to target the RBD, although other potential neutralizing epitopes have also been reported (Cao, et al., 2020; Cheng, et al., 2020; Chi, et al., 2020; Hansen, et al., 2020; Liu, et al., 2020; Pinto, et al., 2020; Renn, et al., 2020; Shi, et al., 2020; Yuan, et al., 2020). The spike is also the protein whose mRNA is used in the two emergency-authorized vaccines.

The UK variant contains up to eight substitutions/deletions in the S protein, including N501Y in the RBD; the South African variant 501.V2 has three (K417N, E484K and N501Y). Understanding the effect of these substitutions on the transmissibility and virulence of SARS-CoV-2 as well as the effectiveness of mAbs known to neutralize the wild-type (wt) S, is of broad and immediate interest.

In the present study, we assess the impact of the three RBD-associated amino acid substitutions in the South African variant 501.V2 spike on the interactions of the RBD with ACE2 and with neutralizing mAbs using a suite of molecular modeling and simulations. Our analysis shows that the mutations K417N and E484K act to counterbalance the high affinity of the spike imparted by the mutation N501Y (shared by both the UK and South African variants), suggesting that the South African variant may be less infectious and less transmissible than the UK variant. Furthermore, most of the structurally characterized mAbs (**Table 1**) including one of the two Regeneron mAbs used in combination (Hansen, et al., 2020) exhibit minimal 3D contacts or compensating interactions at the mutation sites, suggesting that their effectiveness will not be altered, and could even be increased in two cases, B38 (Wu, et al., 2020) and 2-4 (Liu, et al., 2020). In contrast, a weakening in the interactions with the variant is discerned in the cases of mAbs C105 (Barnes, et al., 2020), BD23 (Cao, et al., 2020), REGN10933 (Hansen, et al., 2020) and H11-H4 (Huo, et al., 2020), where the mutations K417N and/or E484K disrupt interfacial interactions that can be only partially compensated by conformational rearrangements. This computational analysis draws attention to the necessity of re-evaluating the neutralizing effect of selected mAbs vis-à-vis the potential ability of South African variant to escape immunotherapy.

**Table 1.**
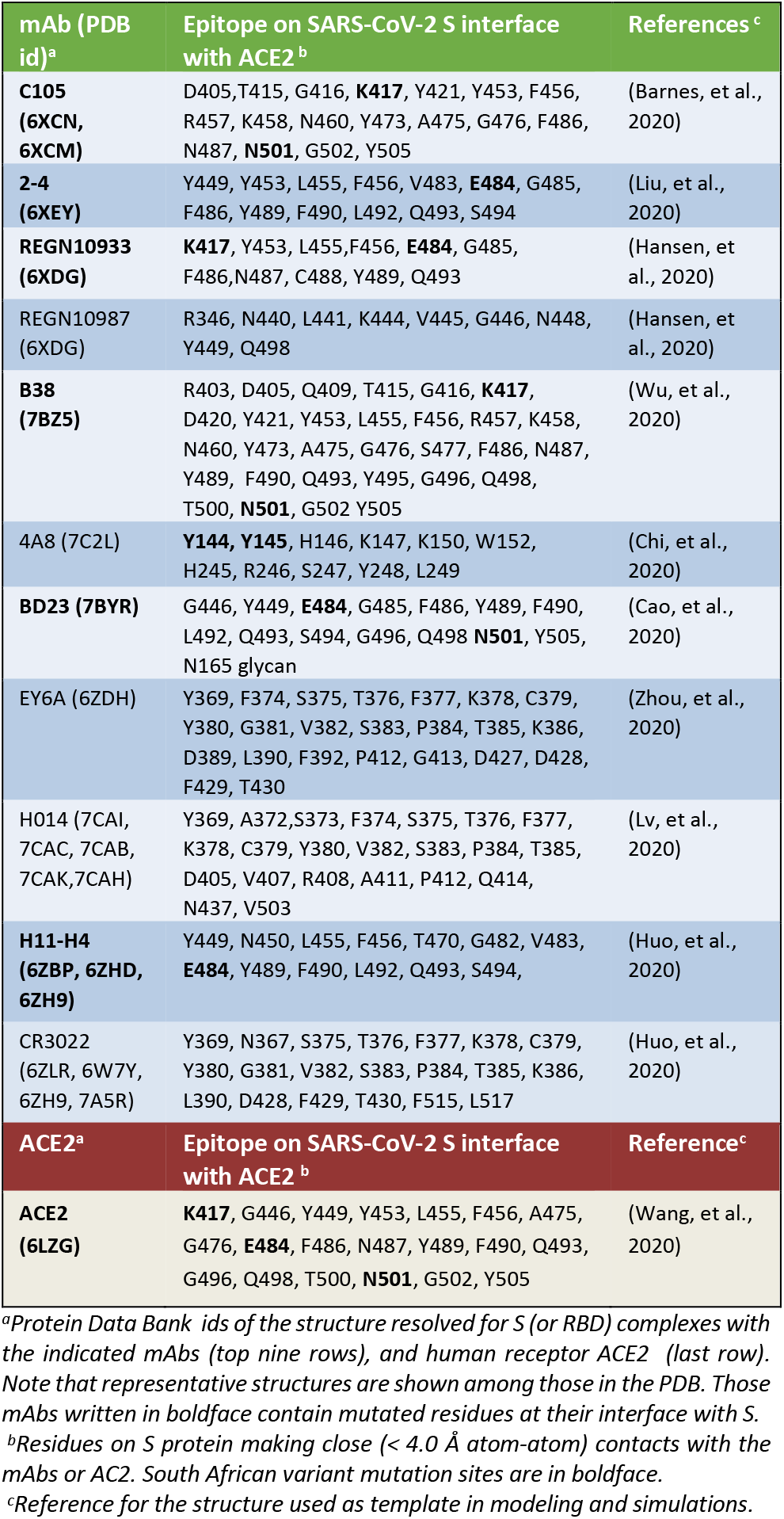
MAbs whose interactions with the South African variant spike have been analyzed in the present study.

## 2 Methods

### *In silico* generation of structural models for SARS-CoV-2 S RBD variants complexed with ACE2 and mAbs

The structure of the complex formed by wt S RBD and ACE2 was obtained from the Protein Data Bank (PDB ID: 6LZG (Wang, et al., 2020)). To characterize the changes in interfacial interactions imparted by the amino acid replacements in the South African 501.V2 variant, we generated structural models using CHARMM-GUI for three mutant RBDs complexed with ACE2: (1) N501Y; (2) K417N and E484; and (3) K417N, E484K, and N501Y. The generated models were energetically minimized and further refined by molecular dynamics (MD) simulations with explicit solvent using HADDOCK 2. 2 (Van Zundert, et al., 2016). Similar modeling and simulations were performed for the complexes of the same variant with a series of mAbs using as template the PDB structures listed in **Table 1**. Those resolved at low resolution (e.g. BD23) were also subjected to energy minimization prior to modeling and simulations.

### Assessment of changes in binding energetics due to amino acid v ariations

For each RBD-ACE2 complex model, four energetically favorable conformations sampled during MD refinement w ere selected to estimate the changes of binding free energy due t o amino acid substitutions, using PRODIGY (Xue, et al., 2016). T he same approach was adopted for the RBD-mAb complexes.

### Characterization of impact of mutations on the structural me-c hanics of the S protein complexed with ACE2

The global dynamics of RBD-ACE2 complex was evaluated using the Gaussian network model (GNM) (Bahar, et al., 1997), as implemented in the *DynOmics* web server (Li, et al., 2017), and by essential site scanning analysis (ESSA) (Kaynak, et al., 2020). Both tools are accessible and described in detail in the open source *ProDy* API (Bakan, et al., 2014; Bakan, et al., 2011). GNM analysis was used to determine the role of the mutation sites in the global dynamics of the S protein (wt or variant) complexed with ACE2, using broadly established analytical methods (Bahar, et al., 2010). Mainly, we evaluated the conformational flexibility (or mean-square fluctuations (MSFs)) of residues in the most cooperative modes of motions and identified the most constrained regions (i.e. minima in these MSFs plotted as a function of residue index, called global mobility profiles). These regions serve as anchors or hinges; as such they play a key mechanical role and cannot easily adapt to substitutions (Haliloglu and Bahar, 2015). ESSA, on the other hand, determine the essential residues whose perturbation alters the frequency dispersion of global motions. Residue-based ESSA scores are evaluated based on three lowest frequency GNM modes. High scores residues usually occupy functional (e.g. orthosteric or allosteric) sites (Kaynak, et al., 2020).

## 3 Results

### N501Y induces an increase in ACE2 binding affinity, and alters the structural mechanics of the RBD-ACE2 complex

The amino acid N501, mutated to Y501 in both the UK and South African variants, forms two hydrogen bonds with the residues Y41 and K353 of ACE2, resulting in a stable complex with ACE2, with a binding affinity of −12.1 ± 0.2 kcal/mol (**Fig. 1B**). N501Y mutation further induces three new associations with ACE2: (i) a hydrogen bond with D38, (ii) a cation-π interaction with K353, and a potentially π-stacking interaction with Y41, resulting in an increase in ACE2-binding affinity to −12.6 ± 0.3 kcal/mol. Note that the interaction partners Y41 and K353 are critical residues that have been reported to regulate ACE2-SARS binding: K353H (A or D) is known to abolish the interaction with the SARS-CoV S glycoprotein; and Y41A strongly inhibits interaction with the SARS-CoV S glycoprotein (Li, et al., 2005). Therefore, in both the UK and South African variants, the substitution N501Y is expected to promote ACE2 binding and thereby induce an increase in transmissibility. This result is consistent with the results from deep mutational scanning experiments (Starr, et al., 2020) as well as a rigorous computational analysis (Fratev, 2020).

**Fig 1.**
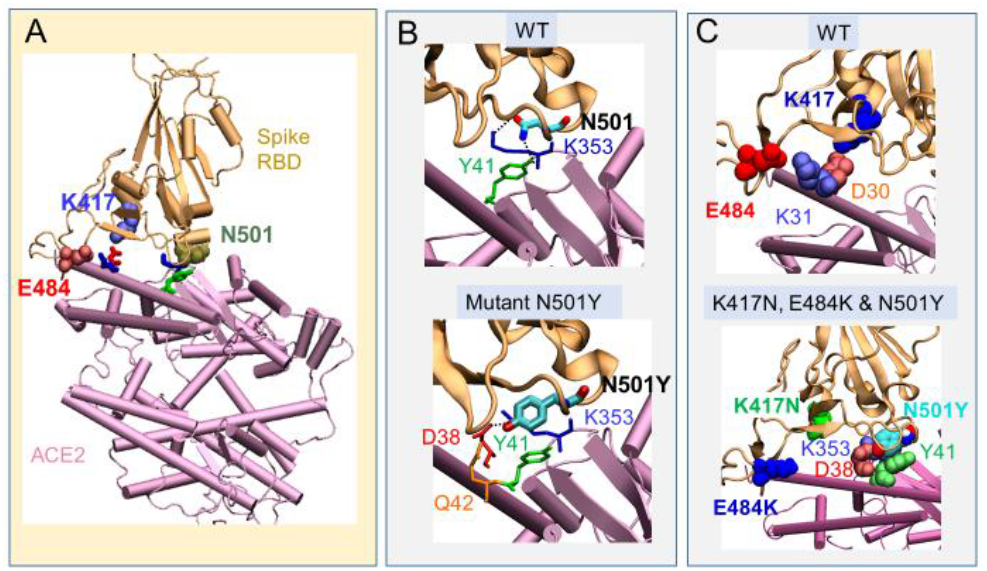
Change in interactions with ACE2 between wt RBD and Southern African variant 501.V2. (**A**) position of the three amino acids (K417, N501 and E584) at the interface with human ACE2 in the WT RBD; (**B**) comparison of the interactions of N501 (*top*) with those of the variant with single substitution N501Y (*bottom*). The latter enables a tighter contact resulting in higher binding affinity compared to WT. (**C**) Same as B for the triple mutant K417N, D501Y and E484K. Two salt bridges originally present in the wt (*top*) are broken (*bottom*), suggesting a compensating effect by the substitutions E484K and K417N, countering N501Y.

### K417N and E484K mutations abolish two salt bridges otherwise formed between ACE2 and S RBD

K417 is specific to SARS-CoV-2 S. The salt bridge between K417 and D30 from ACE2 assists in RBD-ACE2 binding (Wang, et al., 2020). In addition, an interfacial salt-bridge between S E484 and ACE2 K31 further stabilizes the complex (**Fig. 1C** *top*). Notably, the mutations (K417N and E484K) in the South African variant abolish these interfacial salt bridges reducing the binding affinity of the RBD to −11.4 ± 0.3 kcal/mol (in the absence of the 3^rd^ substitution N501Y). This decrease in affinity appears to counterbalance the increase imparted by N501Y, suggesting that the two additional mutations in the South African variant 501.V2 act to compensate the effect of N501Y. This compensating effect is not present in the UK variant.

### Most of the mAbs reported to neutralize the spike show little sensitivity to the substitutions in the South African triple variant

We analyzed a set of eleven antibodies (**Table 1**, first column) whose structures complexed with the SARS-CoV-2 spike or RBD have been resolved. **Fig. 2** displays their binding poses, generated by structurally aligning the spike or RBD regions of the corresponding S/RBD – mAb complex structures. **Table 1** *column 2* lists the residues that make interfacial contacts with the mAbs. Examination of the structures showed that the three mutation sites (K417N, E484K and N501Y) on the South African variant do not engage in direct contacts with five of the listed mAbs: REGN10987, EY6A, HO14, 4A8, and CR3022 (**Suppl Fig 1**), suggesting that their substitutions would affect the effectiveness of these mAbs only minimally, if at all.

**Figure 2.**
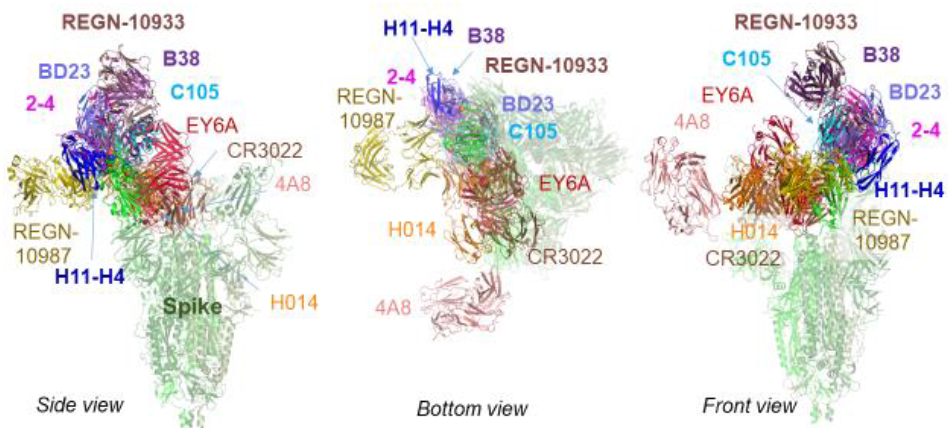
Interactions of the SARS-CoV-2 S with mAbs. See **Table 1** for the list of mAbs and binding epitopes on the S protein.

The remaining six mAbs are engaged in direct contacts with the variant spike at one or more of the three mutation sites. Among them, B38 and 2-4 tend to even exhibit an increased affinity to bind to the variant, compared to their affinity to bind the wt RBD. The corresponding interfacial contacts are illustrated in **Suppl Fig 2A-B**. Mainly, the substitution E484K allows for new favorable interactions with mAb 2-4 residues S54 and N52; and the mutation N501Y leads to a hydrogen bond with mAb 2-4 T28, both in favor of tighter binding of 2-4 to the variant (**Fig 2A** *right panel*). Likewise, K417N creates a new hydrogen bond with B38 residue Y33, and the network of interfacial interactions between RBD pair of residues N501 and Q498 and the mAb B38 pair S67 and S30 is maintained if not strengthened upon substitution of a tyrosine for N501. Quantitative assessment of binding affinities in these two cases show a change of 0.4 – 0.5 kcal/mol in favor of binding, which is small but robustly reproduced in four independent runs.

### The mutations K417N and/or E484K in the South African variant decrease the ability of the mAbs C105, BD23, REGN10933, and H11_H4 to bind the RBD

C105 exhibits a decrease in binding affinity to the variant, compared to its affinity to bind the wt RBD. C105 depends on K417 to coordinate a polar network that supports its high affinity to bind the S RBD. K417 forms a salt bridge with the C105 residues E96/E99 which also engages the spike residue Y453 (see **Fig 3A**). This network is completely abolished by the substitution K417N (highlighted by the *red circle* in panels **A** and **B** of **Fig 3**). This significant loss in the attractive interaction with the mAb C105 is partially alleviated by a new favorable interaction between the N417 and mAb Y52, and a cation-π interaction that Y501 makes with mAb K31. But the net effect is a reduction in binding affinity by 0.4 kcal/mol.

**Figure 3:**
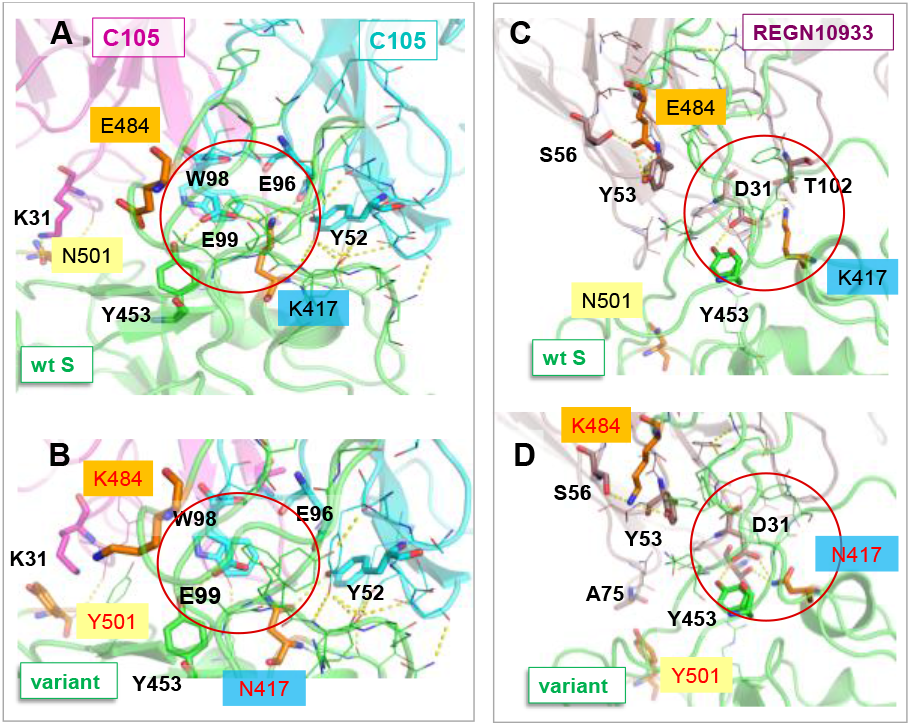
Disruption of salt bridges formed by K417 may weaken the association of C105 and REGN10933 with the South African variant 501.v2, compared to their interaction with the wt S. The panels compare the interactions of the mAbs C105 (A-B) and REGN10933 (C-D) with the wt RBD (A and C) and the South African mutant (B and D). A central salt bridge with C105 (K417-E99/E96) and another with REGN10933 (K417-D31) are lost due to the substitution K417N. A new cation-π interaction with K31 is formed upon N501Y mutation in D, which may compensate to restore the binding of REGN10933. N501Y makes no interfacial contacts, and E484K undergoes rearrangements to alleviate the effect of charge change. The net effect due to those changes in interfacial interactions is a weakening in binding affinity by 0.4±0.2 kcal/mol.

The South African variant spike also appears to engage in a weaker association, compared to the wt S, with three other mAbs: BD23, H11_H4, and REGN10933. **Fig 3C-D** illustrates the change in interfacial interactions in the case of REGN10933, and **Supplementary Fig 3A** and **B** the results for BD23 and H11_H4, respectively. In the wt S-REGN10933 complex, K417 forms a salt bridge with mAb D31 also engaging T102, which in the mutant is replaced by weaker polar interactions. Likewise, E484 forms a salt bridge with BD23 R107 (**Suppl Fig 3A** *middle*), which, on the contrary, gives rise to repulsion between two positively charged groups upon the substitution E484K in the variant. A similar, even stronger effect is observed at the interface between H11-4 and the RBD (**Supp Fig 3B**) where the strong attractive interaction of wt E484 with H11-4 R52 turns into a strong repulsion between R52 and K484.

Overall, the two substitutions K417N and E484K in the South African variant impact salt bridges otherwise formed between the wt RBD and those four mAbs; and the accompanying structural rearrangements do not provide equally strong (new) associations. The net effect is a reduction in the binding affinity of these mAbs by 0.3 to 0.6 kcal/mol. Although this difference is small, it is robustly reproduced in four independent runs for each complex, and it maps to more than 2-fold decrease in the binding affinity of these antibodies, which warrants experimental inspection of dosage-dependent neutralization abilities of these mAbs.

### N501 and K417 are essential residues that mediate the global dynamics of the RBD-ACE2 complex

Apart from interfacial interactions, it is of interest to examine the role of mutated residues in the structural dynamics of the complex formed with ACE2. GNM (Bahar, et al., 1997) analysis of the global dynamics of the RBD-ACE2 complex revealed that N501 plays a central mechanical role, participating in a hinge center (minimum in the global mobility profile) as illustrated in **Fig 4A and D**. Further analysis of essential sites by ESSA (Kaynak, et al., 2020) pointed to two of the residues mutated in the South African variant (K417 and N501; *in red*), and ACE2 two residues (Y41 and K353*; in blue*) that have been reported (Li, et al., 2005) to regulate the ACE2-spike binding (**Fig 4B**). It is interesting to note that several residues gain functional importance (emerge as peaks) upon complexation of the spike with ACE2 (**Fig 4C**), including K417 and N501 in RBD, Y41 and K353 in ACE2, in addition to D30 and K31 in ACE2 located at the interface. We also identified a pocket in the immediate vicinity of K417 and N501 at the RBD-ACE2 interface (**Fig 4E**) which is predicted by ESSA to potentially serve as an allosteric modulator binding site that might interfere with the conformational dynamics of the complex.

**Figure 4.**
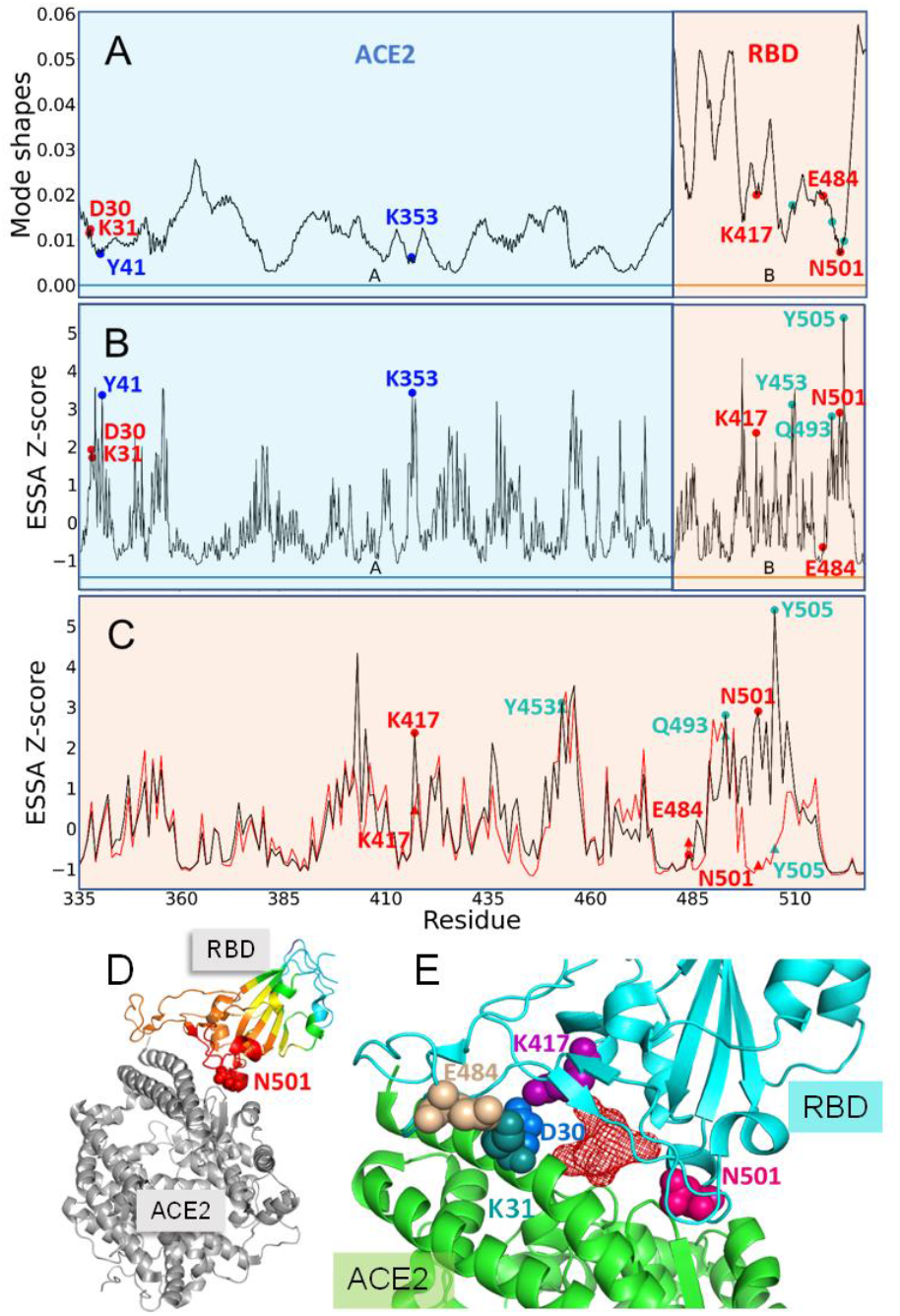
Role of mutation sites and other key sites in the structural dynamics of RBD complexed with ACE2. **A**. GNM-predicted global mobility profiles. Minima (sites participating in global hinge) include RBD N501 and ACE2 Y41 and E353. **B** Results from ESSA of RBD-ACE2 complex. The mutation sites K417 and N501, and associated ACE2 residues (D30 and K31, forming salt bridges with K417 and E484, respectively) are distinguished as peaks (essential residues). Three RBD residues implicated in the human-to-animal passage of the virus are labeled in *cyan*, which are also detected as essential residues. **C.** ESSA profile for isolated RBD (*red curve*), compared to its counterpart in the complex (*black*), shows that N501, K417, and Y505 acquire a significant mechanical role upon complexation. (**D**) Position of N501 in the RBD-ACE2 complex. RBD is color-coded by GNM mobility profile (from *red*; most constrained, to *cyan*, most mobile). (**E**) Closeup view of the interfacial regions displaying an allosteric pocket predicted by ESSA (*red wire)*, and key residues in the vicinity.

## Discussion

SARS-CoV-2 has shown significant mutations in recent months, which raised the question whether any of the new variants would show increased transmissibility, potential to evade monoclonal antibody treatment or altered response to vaccines designed based on wt spike. The high mutation rate may in part be responsible for the zoonotic nature of these viruses and points to a clear risk of still-undetected additional members of the coronavirideae family of viruses making the jump from their traditional hosts to humans in the future.

To date there has been a relatively limited evidence of SARS-CoV-2 mutations that have a significant functional effect on the virus or on the disease. In particular, significant attention has been given to the D614G mutation in the S protein, which emerged early in the pandemic, and might be associated with increased transmissibility (Korber, et al., 2020; Plante, et al., 2020; Volz, et al., 2020; Yurkovetskiy, et al., 2020), and with escape from monoclonal antibody (mAb) and polyclonal serum mediated neutralization (Thomson, et al., 2020). A recent extensive study of 184 recurrent mutations (including D614G) identified in SARS-CoV-2 population (dataset of 46,723 SARS-CoV-2 isolates, sourced July 30, 2020) showed, however, no evidence of increased transmissibility (van Dorp, et al., 2020).

Yet, in more recent months, the emergence of new UK and South African variants, which share a mutation N501Y at a critical (ACE2-binding) site, have once again raised concerns of high transmissibility. Furthermore, mutations in the S protein could affect the efficacy of neutralizing Abs. Indeed, missense mutations that arose in MERS and SARS-CoV resulted in resistance to neutralizing antibodies for the original strains (Rockx, et al., 2010). One hypothesis for the emergence of immune escape variants, is that they arise through intra-host evolution in individuals with immunodeficiency and prolonged viral replication (Avanzato, et al., 2020; Choi, et al., 2020). Indeed, the N501Y mutation is one of the several spike mutations that emerged in an immunocompromised individual in the US who had prolonged viral replication for over 20 weeks (Choi, et al., 2020).

In the present study, we focused on the South African 501.V2 variant and conducted a structure-based computational assessment of the effect of the corresponding substitutions N501Y, K417N and E484K in the S protein RBD on the interactions with ACE2 and neutralizing mAbs using the wealth of structural data accumulated to date for various complexes of the S protein or its RBD with human ACE2 and many mAbs. While the full biological and clinical implications of the new variants in the UK and South Africa are yet to be determined, our study suggests that the affinity of the variant spike to recognize and bind to human ACE2, which is enhanced by the mutation N501Y, but weakened by the interactions engaging the other two mutation sites. As a result, current analysis suggests that despite the stronger binding to ACE2 caused by the substitution N501Y also noted in earlier experiments(Starr, et al., 2020) and computations (Fratev, 2020) for the point mutation, the South African 501.V2 variant that has undergone two additional mutations (K417N and E484K) is unlikely to exhibit an increased infectivity and likely transmissibility originating from an alteration of its interaction with human ACE2.

The effect of the mutations on existing mAbs is more heterogeneous. Here, we examined 11 mAbs (**Table 1**) reported to have neutralizing effects on wt virus, including two Abs used in the Regeneron combination Ab therapy that has been granted emergency use authorization. Five mAb binding epitopes (Epitope I to V) on the SARS-CoV-2 RBD have been identified, residues E484 and K417 contributing to Epitopes I and II (Xiang, et al., 2020). These two charged residues play an essential role in binding many mAbs by forming interfacial salt bridges with oppositely charged spike residues. The interactions between spike and four of the mAbs analyzed—C105, BD23, REGN10933 and H11-14, appear to be impacted due to disruption of salt bridges supported by E484 and K417. Specifically, K417 forms salt bridges with C105 E96/E99, and with REGN10933 D31; and E484 forms a salt bridge with BD23 R107 and with H11-14 R52. These attractive interactions are broken by the variant mutations, and in the case of the E484K mutation, may even be replaced by repulsions. Our analysis after considering local rearrangements that partially offset these adverse effects suggests that the RBD-binding probability of C105, BD23 and H11-14 may be reduced by a factor of two, while the effect on REGN10933 may be less than 2-fold.

On the contrary, for five of the examined mAbs, the mutations are predicted to exert only minimal effects on their interactions with RBD. It was also interesting to note that Fab 2-4 may have gained a 2-fold increase in its binding affinity, as E484K may promote an interfacial cation-π interaction.

A recent mapping of RBD mutations that escape REGN-COV2 cocktail and Eli Lilly’s LY-CoV016 antibodies (Starr, et al., 2020), using a deep mutational scanning method (Greaney, et al., 2020) revealed several sites whose point mutations mediate escape. K417 is indeed observed among those sites of ‘strong escape’ for REGN10933 (but not REGN10987) consistent with our study. Interestingly the mutation K417N is also the ‘top’ site of escape for LY-CoV016. The study further revealed that F486 is the strongest such site that escaped neutralization by REGN10933, and K444-G446 escaped neutralization by REGN10987. We note that these residues make direct contacts with the respective mAbs (**Table 1**). But interestingly the mutation (E406W) escaped the cocktail of both REGN antibodies even though it does not make direct contacts with either mAb in the resolved structure, inviting attention to the necessity of considering structural changes induced by such drastic substitutions, which may not be revealed by existing structures of the complex between the wt spike and mAbs.

Overall, our analysis suggests that a loss in mAb efficacy by the mutations may be alleviated by combination treatments consisting of mAbs that bind alternative, non-overlapping epitopes and act via different neutralization mechanisms to block virus mutational escape (Baum, et al., 2020; Hansen, et al., 2020; Marovich, et al., 2020; Xiang, et al., 2020). A potential approach to minimize the impact of mAb escape mutations is indeed to develop additional mAbs with epitopes that are highly resistant to viral escape, such as those that include epitopes outside of the RBD and epitopes that are cross-reactive across SARS-CoV and SARS-CoV-2, indicating conserved epitopes with low tolerance for mutation (Pinto, et al., 2020; Wec, et al., 2020). We have recently described such a mAb (Cheng, et al., 2020) that binds the putative superantigenic-like motif of SARS-CoV-2 spike (Cheng, et al., 2020). That particular mAb, 6D3, presents the dual advantage of potentially blocking the S1/S2 cleavage site thus interfering with proteolytic cleavage that is essential for viral entry, in addition to targeting the superantigenic region that may contribute to multisystem inflammatory syndromes in in adults and in children and adolescents with severe COVID-19 (Cheng, et al., 2020).

In summary, we provide a computational analysis of the effect of the triple mutations in the South African variant on the infectivity and on the effectiveness of mAbs. The computational predictions provide testable hypotheses, such as that the optimal dosage in selected mAb treatments which may be affected by the mutations. Given the growing access to therapeutic mAbs via clinical trials and emergency use authorization, and the potential emergence of immune evasion mutations that maintain virulence or those that confer resistance to immunizations or therapies (Weisblum, et al., 2020), this type of *in silico* assisted genomic/molecular surveillance may provide feedback for accelerating the design of experimental studies in response to the pandemic.

## Acknowledgements

We gratefully acknowledge support from the NIH awards 3RO1AI072726-10S1 (to MA) and P41GM103712 (to IB) and a MolSSI COVID-19 Seed Software Fellowship (to JK).

## Conflict of Interest

none declared.

## Notes

### Competing Interest Statement

The authors have declared no competing interest.

