## Supplemental figures for "Impact of South African 501.V2 Variant on SARS-CoV-2 Spike Infectivity and Neutralization: A Structure-based Computational Assessment"

*Structural Bioinformatics*

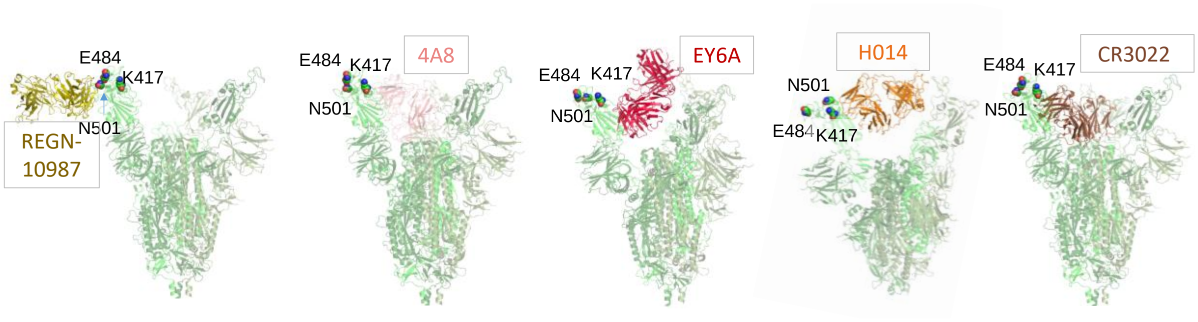


**Figure S1**: **Binding pose of five mAbs that do not make interfacial contacts with the spike at the mutation sites of the South African variant.** Monoclonal antibody (mAb) REGN10987 (Hansen, et al., 2020), 4A8 (Chi, et al., 2020), EY6A (Zhou, et al., 2020), H014 (Lv, et al., 2020) and CR3022 (Huo, et al., 2020; Yuan, et al., 2020) are not affected by the three RBD mutations in the South African mutant. The mutated sites in the RBD are not within the binding epitopes of these mAbs. Structural analysis was performed on the crystal structures of mAbs in complexes with SARS-CoV-2 Spike or RBD. The PDB IDs for the binding of mAb REGN10987, 4A8, EY6A, H014 and CR3022 are 6XDG, 7C2L, 6ZDH, 7CAI and 6ZH9, respectively. The mAb-Spike pose was plotted either directly from crystal structures (with Spike) or after aligning the resolved mAb-RBD complex structure with that of the WT Spike model (PDB: 6VSB) (Wrapp, et al., 2020).

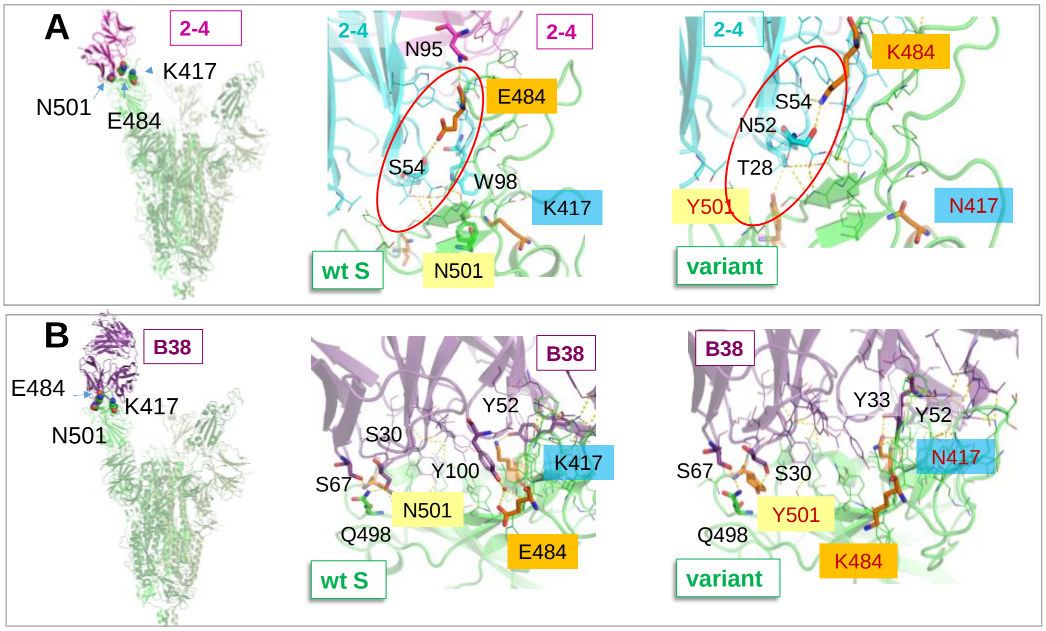


**Figure S2**: **RBD mutations in the South African variant may increase mAb binding affinity for mAb 2-4 and B38** . *Top* and *bottom* panels are for mAb 2-4 (Liu, et al., 2020) (**A**) and B38 (Wu, et al., 2020) (**B**). For each mAb*, left, middle* and *right* panels show mAb bound to the WT Spike, close view of the interfacial interactions of the mAb with the wt RBD, and the South African mutant, respectively. In the case of mAb 2-4 (*top* panels), in the WT spike, E484 makes a hydrogen bond with S54; mutation E484K induces an increase in associations with S54 and N52. In addition, N501Y leads to additional hydrogen bond with T28. As to mAb B38, K417 makes a hydrogen bond with Y52, which is maintained upon mutation to Asn. Furthermore, K417N forms additional hydrogen bonds with Y33, further stabilising the network. N501 or N501Y maintains two hydrogen bonds with S67 and S30. E484 is on the periphery and minorly loses interactions upon mutation to Lys. Overall, the net effects due to mutations may increase the binding affinity of 2-4 and B38 by 0.4±0.2 kcal/mol.

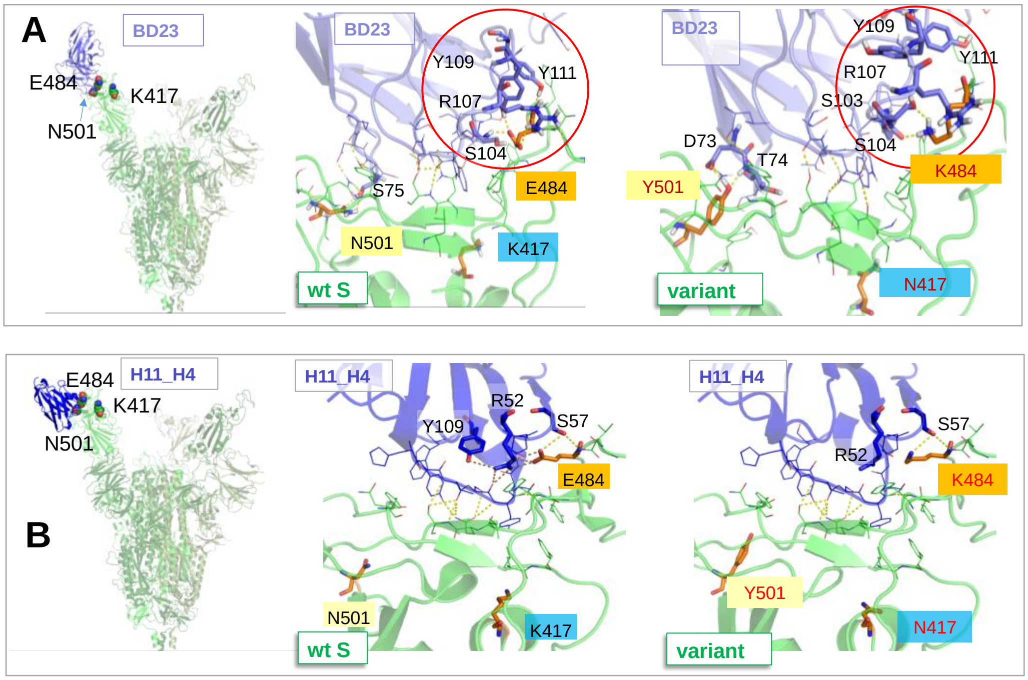


**Figure S3**: **E484K mutation abolishes interfacial salt bridges between mAb and RBD, which may reduce the binding affinity of mAb BD23 and H11_H4.** *Top* and *bottom* panels are for mAb BD23-Fab (Cao, et al., 2020) (**A**) and H11_H4 (Huo, et al., 2020) (**B**). For each mAb*, left, middle* and *right* panels show mAb bound to the WT Spike, close view of the interactions of the mAb with the wt RBD, and the South African mutant, respectively. For BD23 (*top* panels), a salt bridge E484-R107 is diminished due to E484K mutation. Modeling was done based on mAb BD23-bound Spike (PDB: 7BYR). For H11_H4 (*bottom panels*), a central salt bridge E484-R52 is lost due to the E484K mutation, which may induce the escape of the mAb. Note that the other two mutations N501Y and K417N have minor effects on the binding of these two mAbs. Modeling was done based on the H11-H4 bound RBD complex (PDB: 6ZH9). The mAb-Spike pose in (**B**) was plotted by aligning the mAb-RBD complex with the WT Spike model (PDB: 6VSB). For both mAb BD23 and H11-H4, the binding affinity was reduced by 0.6±0.2 Kcal/mol.
